# A Biophysical Model of Phagocytic Cup Dynamics: The Effect of Membrane Tension

**DOI:** 10.1101/2025.11.18.688812

**Authors:** Peyman Shadmani, Behzad Mehrafrooz, Abbas Montazeri, David M. Richards

## Abstract

Phagocytosis is a fundamental cellular process by which cells engulf external particles, controlled by receptor–ligand binding and actin-driven membrane dynamics. While a number of mathematical models have been developed to describe this process, they often overlook membrane tension, a key physical parameter known to influence membrane deformation and cytoskeletal behaviour. To address this gap, we present an enhanced mathematical model of receptor motion during phagocytosis that explicitly incorporates the role of membrane tension. Further, we introduce a signalling component that is coupled to receptor dynamics via the membrane tension. We find that including tension results in fundamentally different engulfment behaviour, which is slower than that predicted by models without tension. In particular, unlike in the previous version of this model, we show that tension can lead to stalled engulfment, an experimentally-observed phenomenon known as frustrated phagocytosis. We also find that signalling is able to modify engulfment behaviour, especially at later stages, and is able to alter cup growth to become linear in time without the need for receptor drift as introduced in previous models. These findings offer new insights into the role of membrane tension and biophysical regulation in phagocytosis, with implications for immune function, cell motility and targeted drug delivery.

## Introduction

Phagocytosis belongs to group of processes called endocytosis, which involves the active transport of cargo into the cell [1]. In particular, phagocytosis is a receptor-mediated, actin-dependent form of endocytosis that is responsible for ingesting relatively large objects, typically over 500nm [2]. It has a variety of uses including removing unwanted cells and debris, and as a way of acquiring nutrients.

Within the immune system, phagocytosis is a critical mechanism needed to eliminate pathogens and initiate other immune responses. It operates in both the innate immune system (which provides a rapid, non-specific defence) and the adaptive immune system (which is slower but antigen-specific and capable of immunological memory) [3]. Phagocytosis within the immune system is performed by specialised cells known as professional phagocytes, which include neutrophils, macrophages, mast cells and dendritic cells [4]. These detect infection sites, engulf foreign particles and present antigens to lymphocytes, thereby activating the adaptive immune response and triggering inflammation. Non-professional phagocytes like epithelial cells, fibroblasts and endothelial cells also perform phagocytosis, though with limited efficiency due to fewer pathogen-recognition receptors [5, 6].

Phagocytosis is a complex, multi-step process. The initial stage, recognition, is mediated by a diverse set of membrane-bound receptors that bind target ligands either directly or via intermediary molecules known as opsonins (including antibodies and complement proteins) [7]. Once receptors bind their ligands, they trigger intracellular signalling cascades involving kinases, adaptor proteins and small GTPases. This signalling promotes receptor clustering and cytoskeletal rearrangement, initiating the formation of the phagocytic cup. Actin polymerisation at the site of target engagement drives membrane extension around the target, and this actin-driven force is essential for successful engulfment, especially of large targets [8]. Motor proteins such as myosins also contribute to the final closure of the phagocytic cup [9].

Mathematical modelling offers the opportunity to understand complex processes throughout biology, not least phagocytosis, which involves diverse biophysical and biological processes such as receptor motion, actomyosin dynamics, intracellular signalling and membrane remodelling. Over recent decades, several models focusing on different aspects of phagocytosis have been developed [10–12]. One of the first was by Petri *et al*. in 1987 who proposed a phenomenological model treating phagocytosis as an irreversible bimolecular reaction, with phagocytes acting like macromolecules containing multiple binding sites [13]. The model quantified the rate and capacity of phagocytosis using a rate constant and maximum uptake per cell.

Since then, increasingly sophisticated models have emerged, utilising a variety of approaches, including finite element modelling [14, 15], Monte Carlo simulations [16] and the finite difference method [17]. In particular, Herant *et al*. proposed a finite element model to evaluate the membrane tension during phagocytosis, finding that tension remains low during engulfment due to membrane insertion from internal stores [14, 15]. They also showed that phagocytosis involves a protrusive force driven by repulsion between the cytoskeleton and free membrane, and a flattening force from cytoskeletal attachment at the leading edge, likely mediated by unconventional myosins. van Zon *et al*. developed one of the first models to incorporate receptor motion during phagocytosis, highlighting how variations in receptor density, membrane curvature and actin recruitment can lead to distinct engulfment outcomes [18]. By identifying a mechanical bottleneck at half-engulfment, their work explains the experimentally-observed bimodal distribution of cup progression, either stalling before halfway or completing entirely. Tollis *et al*. instead used a Monte Carlo approach to elucidate the role of the cytoskeleton in phagocytosis, demonstrating that actin-driven force generation is not strictly required for engulfment [16]. Their model showed that passive receptor-ligand binding can drive phagocytic cup formation and successful uptake of small particles, though the process is slower and more variable. These predictions were supported experimentally using fibroblasts expressing mutant Fc*γ* receptors incapable of actin signalling.

Recent observations suggest that phagocytosis is not a continuous processes, but a set of individual punctuated events. In other words, phagocytosis occurs via a number of distinct stages [19, 20]. For example, Kress *et al*. have shown that filopodia within macrophages retract by discrete steps [21]. The first mathematical model to account for this discrete nature of phagocytosis was proposed by Richards and Endres [17]. In this model, the motion of receptors within the membrane is mapped to the well-known Stefan problem of phase transitions. This was later extended to a three-dimensional model that allowed the effect of target shape to be investigated [22].

Shadmani *et al*. developed an energy-based model that included a role for the plasma membrane tension [23]. They investigated the impact of a protein corona on receptor-mediated endocytosis and their simulations showed that membrane tension, while negligible for small particles, becomes a major limiting factor for larger ones. This model was able to elucidate the roles of various energy contributions in either inhibiting or facilitating receptor-mediated endocytosis.

Recently, Gov *et al*. developed a simplified coarse-grained model for simulating phagocytosis [24]. Their model accounts for various contributions to the energy, including the bending of the cell membrane, adhesion between the target and cell, actin polymerisation and chemical binding energy. They employed the Metropolis algorithm to minimise the system energy and predict its subsequent state over time. Utilising their model, they explored the impact of target size, shape and orientation, showing that spherical particles are engulfed more easily than non-spherical ones of the same surface area, and that the orientation of non-spherical targets strongly influences engulfment dynamics. They also demonstrated that curved membrane-bound proteins self-organise at the leading edge of the phagocytic cup, reducing the bending energy and facilitating faster internalisation. They validated their model through experimental data, specifically examining the phagocytosis of polystyrene beads by macrophages.

The size and shape of the target has been shown to have a dramatic effect on the outcome of phagocytosis [25–28]. Among the early work to investigate the role of target size is that of Tabata and Ikada [26] and Rudt *et al*. [28], who found that phagocytosis is maximised for microspheres in the size range of 1-2 *µ*m. Rudt *et al*. further showed that surface coatings with long-chain poloxamers could nearly eliminate phagocytosis by sterically stabilising the particle surface. A similar study was conducted by Simon *et al*., who reported that increased target size leads to greater demand on the cell’s membrane reservoir, ultimately limiting uptake once the available membrane area is exhausted [27].

These studies were performed using spherical targets. However, the role of target shape has also been studied by various groups. These include Champion and Mitragotri, who showed that particle shape, not size, is the dominant factor in determining whether phagocytosis is initiated [29]. Specifically, they found that local geometry at the point of contact determines whether actin structures can form to support engulfment, with highly curved tips facilitating uptake more readily than flatter regions. Doshi and Mitragotri also determined that macrophage attachment is strongly influenced by particle geometry, with the greatest recognition occurring for shapes with a longest dimension of 2–3 *µ*m, matching the size of many bacteria [30]. More recent work by Gov *et al*. [24] and Richards and Endres [22] showed that phagocytosis of non-spherical targets is highly sensitive to particle orientation, and that tip-first presentation often results in faster or more successful engulfment. Paul *et al*. also demonstrated experimentally that ellipsoidal particles are engulfed significantly more slowly [31].

In this paper, based on observations that membrane tension plays an important role during phagocytic engulfment [14, 23, 24, 32], we extend the model in Richards *et al*. by incorporating the role of membrane tension [17]. We also consider an improved form of signalling, with a more realistic link between signalling and engulfment. Using our improved model, we then investigate the dynamics of engulfment, with a particular focus on how variations in cellular mechanical properties (such as membrane tension and bending stiffness) and target properties (such as size and ligand density) influence phagocytic behaviour.

## Model development

We base our approach on the model of Richards *et al*. [17], which itself is based on the Gao *et al*. model of receptor-mediated endocytosis [33]. This model focuses on the motion of receptors in the cell membrane (see Figure 1). By moving to the edge of the phagocytic cup, receptors are able to bind ligands on the target particle and so progressively increase the size of the bound region until the target is completely engulfed.

**Fig 1.**
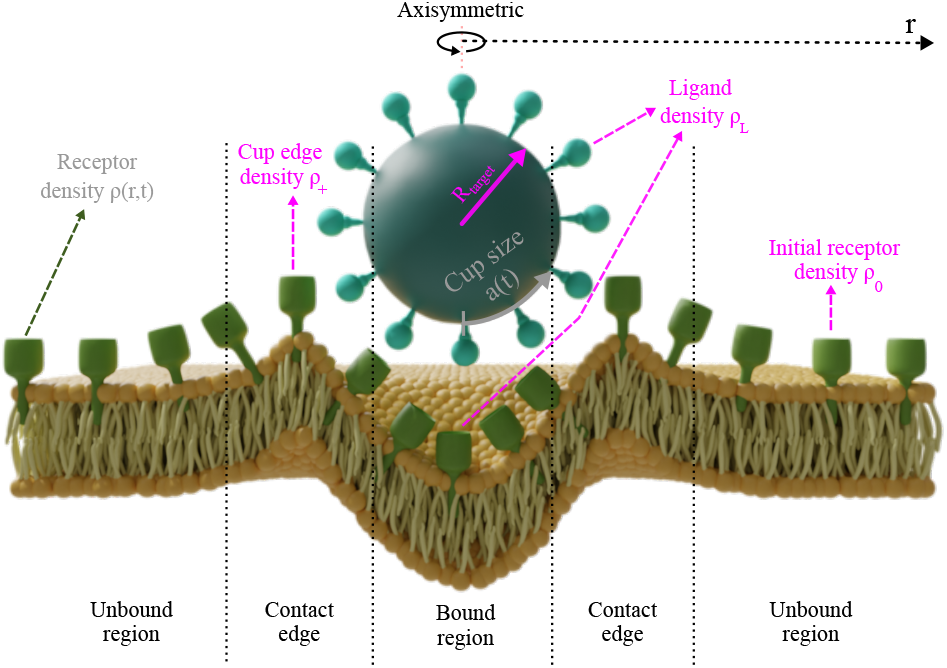
**Model schematic showing the three main regions of the membrane**: the bound region within the phagocytic cup, the contact region just outside the cup (where variables are labelled by the subscript +) and the unbound region where receptors are free to diffuse. The three model variables are the receptor density *ρ*(*r, t*), the signalling molecule density *S*(*r, t*) and the cup size *a*(*t*). Important other quantities include the initial receptor density *ρ*_0_, the ligand density *ρ*_*L*_ and the receptor density at the edge of the cup *ρ*_+_.

For spherical targets (which is all we consider here) in the absence of noise, the system displays circular symmetry when viewed from directly above the target. This means that motion in the two-dimensional membrane only depends on the radial distance *r* away from the cup. Thus the density of receptors, our first model variable, can be described as *ρ*(*r, t*), where *t* is the time. Initially, before any contact between the cell and the target, we assume a constant density of receptors, *ρ*(*r*, 0) = *ρ*_0_.

The second model variable is the phagocytic cup size, *a*(*t*), defined as half of the engulfed arc length. At the beginning of phagocytosis this starts at zero and then gradually increases until full engulfment when *a*(*t*) = *πR*, were *R* is the target radius. We assume that receptors cannot unbind ligands, leading to a zipper-like mechanism where *a*(*t*) can never decrease. This agrees with several experimental results and previous models [17, 18, 34].

Our model also includes a simple role for intracellular signalling. This is a vital part of phagocytosis, which is much more active than other types of endocytosis [35]. Although the signalling pathways are complex [8, 36], involving many different proteins and interactions, for simplicity we consider only a single signalling molecule that resides within the membrane. This is described by our third and final variable, *S*(*r, t*), which represents the density of the signalling molecule at every point along the membrane.

We now describe in detail the two parts of our model—receptor motion and signalling dynamics—along with how these are coupled together.

### Receptor motion

There are a number of possible ways that receptors can move, include passive processes such as diffusion and active processes such as motion along actin filaments. Although previous models have included both of these, we here only consider diffusive behaviour. In fact, one of the main results we find is that, by coupling signalling to receptor motion in the correct way, it is possible to obtain previous engulfment behaviours without the need of also introducing active, drift-like dynamics. This does not mean that active processes are not present during phagocytosis, simply that they do not need to directly influence receptor motion. Rather active processes are likely to influence other properties of the system, such as the membrane tension, membrane curvature or receptor-ligand binding.

Since the cell membrane is two-dimensional (albeit with circular symmetry), we describe the receptor dynamics using the two-dimensional diffusion equation

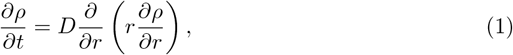

where *D* is the diffusion coefficient. For the boundary condition far from the cup, we assume that the receptor density is always that of the initial density, *i*.*e. ρ*(*r* = ∞, *t*) = *ρ*_0_ for all times.

Upon first attachment of the target to the cell, a localised increase in receptor density occurs within the bound region. We assume this is because the ligand density *ρ*_*L*_ is larger than *ρ*_0_ and that all ligands within the cup are bound to receptors, *i*.*e. ρ*(*r, t*) = *ρ*_*L*_ for *r < a*(*t*) (see Figure 1). Due to this increased density within the cup region, there is a lowered density just outside the cup. The density at the very edge of the cup plays an important role in our model and is represented by *ρ*_+_ ≡ *ρ*(*a*(*t*), *t*). As explained in our previous work, based on the conservation of membrane-bound receptors, the cup growth rate can be expressed as [23]

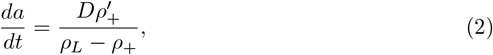

where *ρ*^*′*^_+_ is the derivative of the receptor density with respect to *r* evaluated at the cup edge.

### The receptor density at the cup edge

To have a unique solution, the above system needs one more piece of information, which we take as a the receptor density at the edge of the cup, *ρ*_+_ = *ρ*(*a*(*t*), *t*). This is a moving boundary condition because the cup size is continually growing. As in previous models, we determine *ρ*_+_ by considering the free energy.

We include four contributions to the free energy, with the first three the same as in previous models [22, 23]. First, we consider binding between the cell and the target. With an energy per receptor-ligand bond of −*k*_*B*_*TC*_*b*_, where *C*_*b*_ is the binding constant, *k*_*B*_ the Boltzmann constant and *T* the temperature, the contribution of binding to the total energy is −*πk*_*B*_*Tρ*_*L*_*C*_*b*_*a*(*t*)^2^. Second, based on Helfrich’s classic expression for the membrane bending energy per unit area of 2*k*_*B*_*TC*_*c*_*H*^2^ [37], where *C*_*c*_ is the elastic bending modulus and *H* is the mean curvature, we find a total bending energy of 4*πk*_*B*_*TC*_*c*_*H*^2^*a*(*t*)^2^. For a spherical target (which is all we consider here), *H* = 1*/R*_target_, where *R*_target_ is the target radius. Third, the configurational entropy contributes 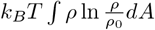 to the free energy, where the area integral extends over the entire membrane [38].

One of the key novelties in this study is the inclusion of an additional, fourth contribution to the free energy, that due to the membrane tension. As phagocytosis proceeds and the cell wraps around the target, there is typically an increase in membrane area with an associated elastic stretching energy. To model this, we consider a membrane of initial area *A*_0_ and tension per unit length of *τ*_0_. When the membrane area is increased by Δ*A*, we assume that the tension per unit length becomes 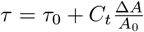, where *C*_*t*_ is the tension coefficient [39]. Then the energy due to membrane stretching is given by [40]

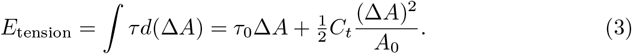

Since the cup area is approximately Δ*A* = 2*πa*(*t*)^2^ (assuming the membrane within the cup bends back on itself as explained in Richards *et al*. [17]) and given that a spherical cell of radius *R*_cell_ has surface area 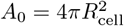, the final energy due to tension is

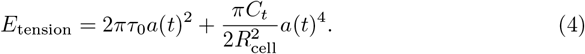

An important point to note is that, unlike the other contributions to the free energy that depend on *a*(*t*)^2^, the tension energy includes a term that depends on *a*(*t*)^4^. This difference is at the root of much of the interesting behaviour that we find below.

Including all four contributions, the system’s total free energy is given by

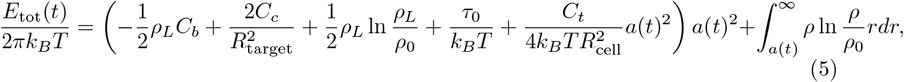

where the *τ*_0_ and *C*_*t*_ terms are the new terms due to membrane tension.

Differentiating *E*_tot_(*t*) with respect to time gives

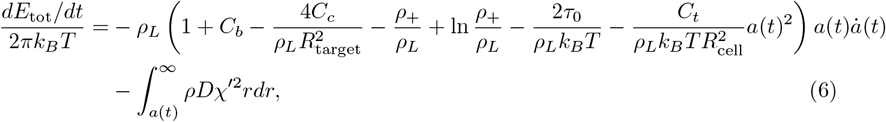

where 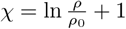 represents the chemical potential per membrane receptor. The integral represents the rate of energy dissipation in the unbound region, so that the remaining terms can be identified as the free-energy jump across the edge of the cup. Assuming this jump vanishes gives an expression for *ρ*_+_, the receptor density at the edge of the cup, as

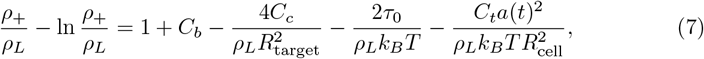

with the final two terms giving the new contribution due to membrane tension.

This last expression gives the boundary condition on *ρ*_+_, the final condition needed to give the system a unique solution. Interestingly, unlike in previous models where *ρ*_+_ was a constant, independent of time, the value of *ρ*_+_ now changes during engulfment as *a*(*t*) gradually increases. Due to this, the system no longer has an analytic solution and a numerical scheme is needed to make progress. Table 1 summarises the initial and boundary conditions.

**Table 1.** Initial and boundary conditions for the system.

| Condition type | Description | Expression |
| --- | --- | --- |
| IC | No engulfment at start | $a(0) = 0$ |
| | Initial uniform receptor distribution | $\rho(r, 0) = \rho_0$ |
| BC | Receptor density far from cup | $\rho(\infty, t) = \rho_0$ |
| | Receptor density at the cup edge | $\rho(a(t), t) = \rho_+$ |

### Signalling dynamics

The second major component of our model is intracellular signalling. Without this, the model is more appropriate to receptor-mediated endocytosis and does not capture the active, energy-dependent aspects of phagocytosis. The various receptors involved in phagocytosis recognise different targets and initiate distinct signalling pathways [41]. These specific signals lead to membrane remodelling and cytoskeletal rearrangement, key aspects of phagocytosis that allow engulfment of targets significantly larger than those possible with other types of endocytosis [42, 43]. A key cellular mechanism during phagocytosis is actin polymerisation, which forms a dense ring of actin filaments around the site where the target attaches to the cell. High levels of actin in this region promote the formation of protrusions on the cell surface, resulting in the formation of a phagocytic cup that progressively wraps around the target.

The full signalling pathways involved in phagocytosis are complex and still to be fully elucidated [35]. In our model, extending the approach by Richards and Endres [17], we abstract all signalling via a proxy membrane-bound signalling molecule, described by its density *S*(*r, t*). As with the receptors, we assume that the signalling molecule only moves via diffusion. However, unlike the receptors, we also allow *S* to degrade with a constant lifetime *η* and to be produced within the cup region at constant rate *βρ*_*L*_ (see Figure 2). This leads to the following model

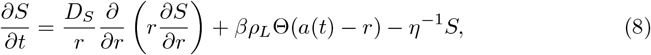

where *D*_*S*_ is the signalling molecule diffusion constant (generally different to the receptor diffusion constant *D*) and Θ is the Heaviside step function, which ensures that *S* is only produced within the cup where *r < a*(*t*).

**Fig 2.**
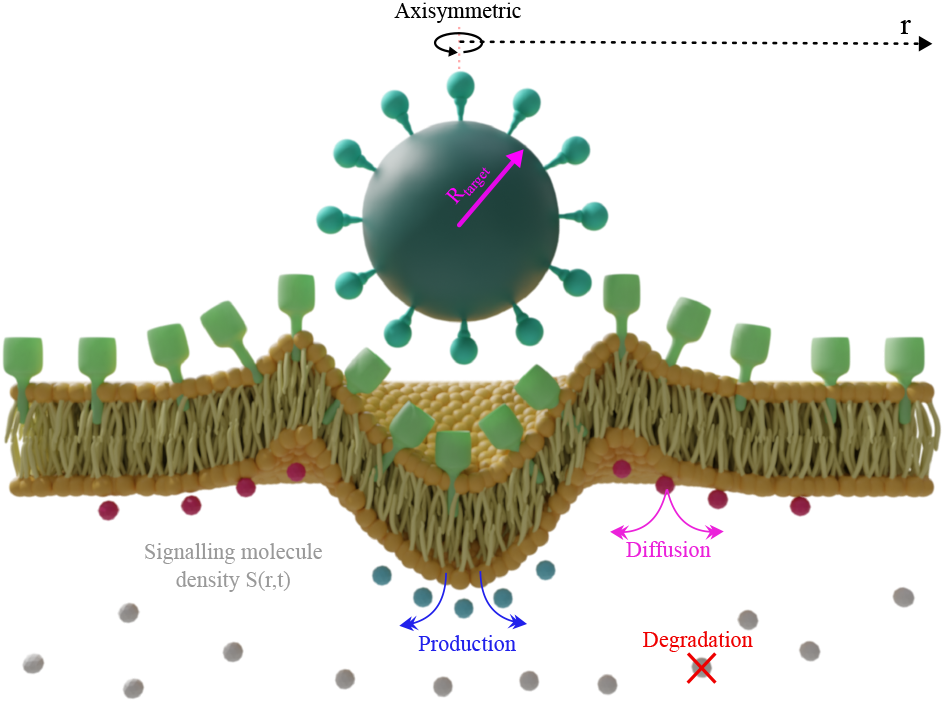
The role of the signalling molecule *S*(*r, t*) in the model. *S* dynamics include three processes: production with the cup region, constant-rate degradation and diffusion.

The last ingredient of our model is how the signalling molecule *S* couples to the receptor density *ρ*. In the previous model by Richards and Endres [17], this coupling was via the receptor drift velocity. Since we do not include drift in our model (and since we do not need to), we instead couple *S* to another part of the system.

In particular, we consider linking *S* with the membrane tension by allowing the tension to reduce with increased signalling. Biologically, this could be achieved either by inserting new membrane (which will decrease the parameter *τ*_0_) or by making the membrane less stiff (which will decrease *C*_*t*_). In the latter case, this could be caused by changing the constituents of the cell membrane itself, but is more likely due to remodelling of the actin cortex directly underneath the membrane. In our model, we capture these effects using either

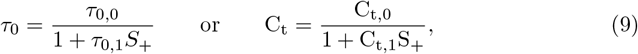

where *S*_+_ is the value of the signalling molecule concentration at the edge of the cup, *i*.*e. S*_+_ = *S*(*a*(*t*), *t*). The new constants *τ*_0,0_ and *C*_*t*,0_ represent the initial values of *τ* and *C*_*t*_ before phagocytosis has begun. As engulfment proceeds and *S*_+_ gradually increases, the membrane tension gradually decreases, with the size of this effect controlled by *τ*_0,1_ and *C*_*t*,1_.

The overall effect of both of these couplings is to gradually increase the right-hand side of Eq. (7) during engulfment. In turn, this decreases the value of *ρ*_+_ and so leads to quicker engulfment than the 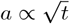 behaviour seen in the no signalling case.

## Results and Discussion

### The effect of membrane tension

We first examine how including membrane tension in our model affects the engulfment dynamics. To understand this fully, we initially consider the model without any signalling so that *τ*_0_ and *C*_*t*_ are constants. In previous models this has been described as a “pure diffusion” model since only the diffusive motion of receptors is considered, without any more active forces such as drift or signalling.

The role of membrane tension in our model is controlled by two parameters: the initial tension per unit length *τ*_0_ and the tension coefficient *C*_*t*_. The effect of *τ*_0_ is simply to add another constant term to the *ρ*_+_ boundary condition (Eq. (7)). In particular, compared to a model that does not include the membrane tension and has all other parameters identical, the inclusion of *τ*_0_ increases the value of *ρ*_+_ and so slows engulfment. Further, a sufficiently large value of *τ*_0_ will lead to *ρ*_+_ *> ρ*_0_ when engulfment is never able to start.

The effect of the tension coefficient *C*_*t*_ is more non-trivial. Due to presence of the *a*(*t*)^2^ multiplying this coefficient in Eq. (7), the value of *ρ*_+_ increases during engulfment (as *a*(*t*) itself increases). This means there is no longer an analytic solution and *a*(*t*) no longer increases with the square-root of time. Instead wrapping of the phagocytic cup becomes slower and slower (compared to 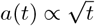) as engulfment proceeds (see Fig. 3A).

**Fig 3.**
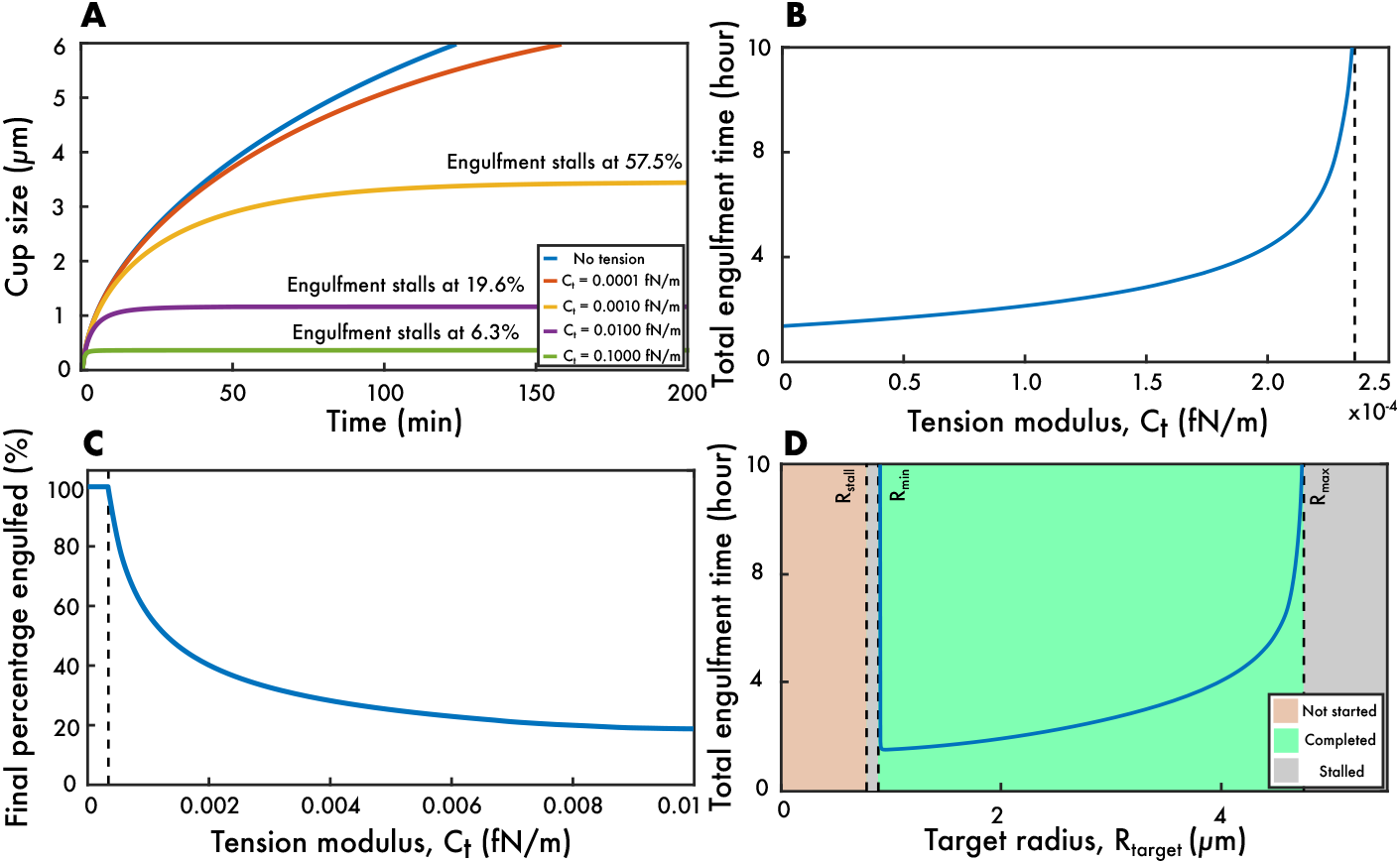
The effect of membrane tension on phagocytosis. (A) Cup size *a*(*t*) against time for various tension coefficients *C*_*t*_. In the absence of tension, the cup grows with the characteristic 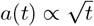 behaviour. However, with tension this is no longer the case and for sufficiently large values of *C*_*t*_ engulfment stalls before completion. (B) Total engulfment time against *C*_*t*_ showing divergence to infinite times as the critical *C*_*t*_ is reached. (C) Maximum percentage of target engulfed against *C*_*t*_. Above the critical *C*_*t*_, engulfment stalls and the percentage engulfed drops below 100%. Increasing *C*_*t*_ further show progressively less and less engulfment, with the largest *C*_*t*_ resulting in barely any engulfment at all. Parameter values for panels (A)-(C): *D* = 1*µ*m^2^*/*s, *C*_*c*_ = 10, *C*_*b*_ = 15, *τ*_0_ = 0.014fN*/*m, *k*_*B*_*T* = 4 *×* 10^−21^J, *ρ*_*L*_ = 500*µ*m^−2^, *ρ*_0_ = 50*µ*m^−2^, *R*_target_ = 2*µ*m, *R*_cell_ = 10*µ*m. (D) The total engulfment time as a function of the target size for *C*_*t*_ = 10^−4^fN*/*m with the other parameter values as above. Sufficiently small targets cannot be completely engulfed due the high bending energy. Similarly, sufficiently large targets stall before complete engulfment due to excessively high membrane tension. Interestingly, there is an intermediate target size corresponding to the shortest engulfment time.

The total engulfment time against *C*_*t*_ is plotted in Fig. 3B, showing that the effect of tension can substantially increase the time for complete internalisation, with larger tension constants leading to longer engulfment times. It is worth noting that all these engulfment times are considerable longer than those observed experimentally. This is because we have not yet included the signalling component to our model; doing so later will result in more realistic total engulfment times.

At sufficiently large values of *C*_*t*_, engulfment stalls before completion (see Fig. 3A). This experimentally-observed phenomenon [44, 45], often called frustrated phagocytosis, has been captured in previous mathematical models [18], but was not seen in our previous receptor-based model. This reveals the existence of a critical value of *C*_*t*_ above which phagocytosis fails to complete. In Fig. 3B this is associated with the total engulfment time tending to infinity.

To investigate this further, we examined the stalling point across a range of membrane tension constants, with the results presented in Fig. 3C. Above the critical *C*_*t*_ value, the maximum percentage engulfed decreases with increasing *C*_*t*_. Although phagocytosis is always able to begin, for very large tension coefficients engulfment stalls when the target is only minimally wrapped.

We next examined the effect of target size *R*_target_ on the total engulfment time. Interestingly, the fate of phagocytosis separates into four regions (see Fig.3D). First, for sufficiently small targets, engulfment can never even begin. This is because such targets are so curved that the cell membrane cannot wrap around them: the associated curvature energy is so large that *ρ*_+_ is above *ρ*_0_ even at *t* = 0.

As the target radius is increased the curvature term decreases and eventually engulfment is able to begin. We call the radius at which this happens *R*_stall_, whose value can be found by solving Eq. (7) with *ρ*_+_ = *ρ*_0_ and *a*(*t*) = 0. This leads to the second region in Fig.3D, where engulfment begins but stalls before completion. This is because, although *ρ*_+_ starts below *ρ*_0_ in such cases, the presence of the *a*(*t*)^2^ term in Eq. (7) means that *ρ*_+_ rises above *ρ*_0_ before the target is completely engulfed.

As the size of target is increased further, eventually a point is reached when phagocytosis completes (the third region in Fig.3D). The minimum target radius for this to occur is labelled *R*_min_. In sharp distinction to our previous model, this region only has finite width. For sufficiently large targets, *i*.*e*. those with radius above *R*_max_, the *a*(*t*)^2^ term becomes so large that engulfment again stalls, leading to the final region in Fig.3D. Thus our model correctly captures the experimental fact that both sufficiently small and large targets can not be phagocytosed.

The precise values of *R*_min_ and *R*_max_ can be found by solving the quadratic equation obtained from Eq. (7) with *ρ*_+_ = *ρ*_0_ and *a*(*t*) = *πR*_target_. It is worth noting that the three critical radii that we identified are related by 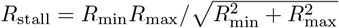 For likely values of *R*_min_ and *R*_max_, this implies that *R*_stall_ is only slightly below *R*_min_, perhaps explaining why frustrated phagocytosis of small targets has not yet been experimentally observed.

Within the region where phagocytosis fully completes (the third region), the engulfment time shows interesting non-monotonic behaviour: there is an optimal target size for which engulfment is quickest. Below this optimal size engulfment is slower due to the high target curvature. Conversely, above the optimal size, although engulfment is quicker, there is a larger target surface area that thus requires more time to be wrapped. This prediction of an optimal target radius for the quickest engulfment agrees with previous models. However, unlike previous models, our model predicts that the optimal target size is only just above *R*_min_, which means the region of decreasing engulfment times is very narrow. Thus experimental studies are likely to miss this region and simply conclude that larger targets take longer to phagocytose.

Our model does not directly take into account the size of the cell and assumes there is infinite membrane available. For a more realistic cell of finite size there will be a target size above which the cell membrane will simply not be large enough to wrap the target. This will modify Fig. 3D to introduce a fifth region at very large target sizes. As with the fourth region, this region will also correspond to stalled engulfment, but now stalling is due to limited available membrane rather than to high membrane tension. The idea of a maximum target size that can be engulfed matches with the ideas of Cannon and Swanson that macrophages have a phagocytic capacity [44]. They observed that as the target size increases macrophages become less effective at phagocytosis, and that beyond a certain size cells are rarely able to complete engulfment.

Within our model we have also ignored the possibility of spare membrane. Cells can often buffer increases in membrane tension through the deployment of membrane reserves (such as membrane folds, microvilli or vesicle fusion) or by creating *de novo* membrane [15, 46]. Although we do not do so here, we could include this in our model by modifying Eq. (4). One way of doing this could be that for small cup sizes (*i*.*e*. small *a*(*t*)) there would be no associated tension energy (*E*_tension_ = 0) as spare membrane is utilised. Only above some particular cup size would the tension energy start to increase.

Finally, it is worth noting that our model treats the cell membrane as an elastic material, where membrane tension is a direct function of surface extension. However, biological membranes also exhibit viscoelastic behaviour and so can at times resist flow and deformation under stress. Potentially this could affect the outcome of phagocytosis. For example, the viscoelastic nature of membranes means that tension build-up during engulfment could be partially relaxed or delayed, allowing the cell to continue wrapping a target even when our simple model predicts stalling [47]. This would effectively postpone or even prevent the onset of frustrated phagocytosis, particularly when engulfing large targets [48].

### Coupling tension to signalling

We now investigate the effect of including signalling alongside receptor motion. As explained above, we do this by coupling the signalling molecule concentration *S* to the membrane tension, either via the initial tension per unit length, *τ*_0_, or via the tension coefficient, *C*_*t*_ (see Eq. (9)).

We focus first on *τ*_0_. Unlike increasing the tension coefficient *C*_*t*_, which slows phagocytosis and can lead to stalling (see Fig.3A), the effect of signalling is to increase the rate of engulfment (see Fig.4A). Not only does signalling decrease the total engulfment time, but it also fundamentally changes how the phagocytic cup grows with time, such that growth is faster than the 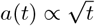 found for the no-signalling, no-tension model. To quantify this growth behaviour, we fit a power law *a*(*t*) = *Bt*^*α*^ to the size of the cup during the first minute of engulfment. Here *α* is the cup growth power and *B* can be understood as the cup size when *t* = 1.

It is notable that signalling can lead to linear cup growth *a*(*t*) ∝ *t* (Fig.3A). This has been reported in experimental work and was only possible in our previous model by modifying the receptor dynamic to include drift in addition to diffusion [17, 22]. Our current model shows that drift is not required to capture linear cup growth, but that the same effect naturally arises by coupling signalling to the membrane tension. In fact, such coupling can even lead to super-linear growth as shown by the *a*(*t*) ∝ *t*^1.4^ growth seen in Fig.3A.

To further understand the effect of signalling on cup growth, we plot *α* against the *τ*_0_-signalling coupling constant *τ*_0,1_ (Fig.3B). As expected, for zero coupling, the cup grows close to the square-root of time (*α* = 0.5). (The growth is not exactly *α* = 0.5 due to a non-zero *C*_*t*_.) As *τ*_0,1_ is increased, the power *α* initially increases. This is because, as the signalling molecule density *S* gradually increases during engulfment, the value of *S* at the cup edge, *S*_+_, increases. This in turn decreases *τ*_0_ (as given by Eq. (9)), which decreases the value of *ρ*_+_ (as given by Eq. (7)), leading to progressively quicker and quicker engulfment.

As *τ*_0,1_ is increased further the power *α* does not increase without limit, but reaches a maximum around *α* = 1.4. Further increasing *τ*_0,1_ gradually reduces *α* back towards 0.5. This is because large values of *τ*_0,1_ result in *τ* increasing very quickly during engulfment, such that *ρ*_+_ quickly decreases towards 0. Once this happens *ρ*_+_ ≈ 0 becomes approximately constant and cup growth becomes 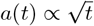 again. Our simulations show that the maximum value of *α* (as a function of *τ*_0,1_) depends on the other parameter values and is likely to have no finite upper limit.

We next examined the total engulfment time as the *τ*_0,1_ coupling is increased (Fig.3C). Unlike in the no-signalling model, which led to unrealistically long engulfment times, our model can now gives engulfment times just over a minute, corresponding well with experimental results. Even though the value of *α* depends non-monotonically on *τ*_0,1_, the total engulfment time gradually decreases as *τ*_0,1_ is increased, tending to a quickest engulfment time when *ρ*_+_ ≈ 0 throughout the whole of engulfment.

We have so far assumed that *τ*_0_ is coupled to signalling via Eq. (9). It is possible to extend this to

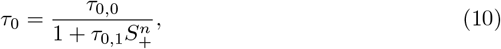

where *n* is a new constant, the Hill coefficient. Increasing *n* above one leads to a sharper transition between high and low *τ*_0_ values as *S*_+_ increases. Sufficiently large values of *n* result in a switch-like transition in *τ*_0_, which was how coupling was implemented in our previous model [17]. The effect of Eq. (10) with large *n* is to split cup growth into two distinct stages. Both stages are well described by 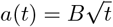 behaviour, but the coefficient *B* is larger in the second, later stage.

Finally, we examine the effect of coupling signalling, not to the initial tension per unit length *τ*_0_, but rather to the tension coefficient *C*_*t*_ (see Fig. 4D). Unlike coupling to *τ*_0_, the cup growth power *α* is now always less than 0.5. This is because the *C*_*t*_ term in Eq. (7) is multiplied by *a*(*t*)^2^ and so is zero at *t* = 0. Thus this term can only increase during phagocytosis and so engulfment cannot proceed quicker than the classic 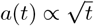 behaviour. For small values of *C*_*t*,1_, engulfment is slowed down as in Fig. 3A, leading to values of *α* under 0.5. The effect of signalling in this case is to reduce the effect of tension at later stages of engulfment, with the largest values of *C*_*t*,1_ restoring 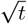 behaviour.

**Fig 4.**
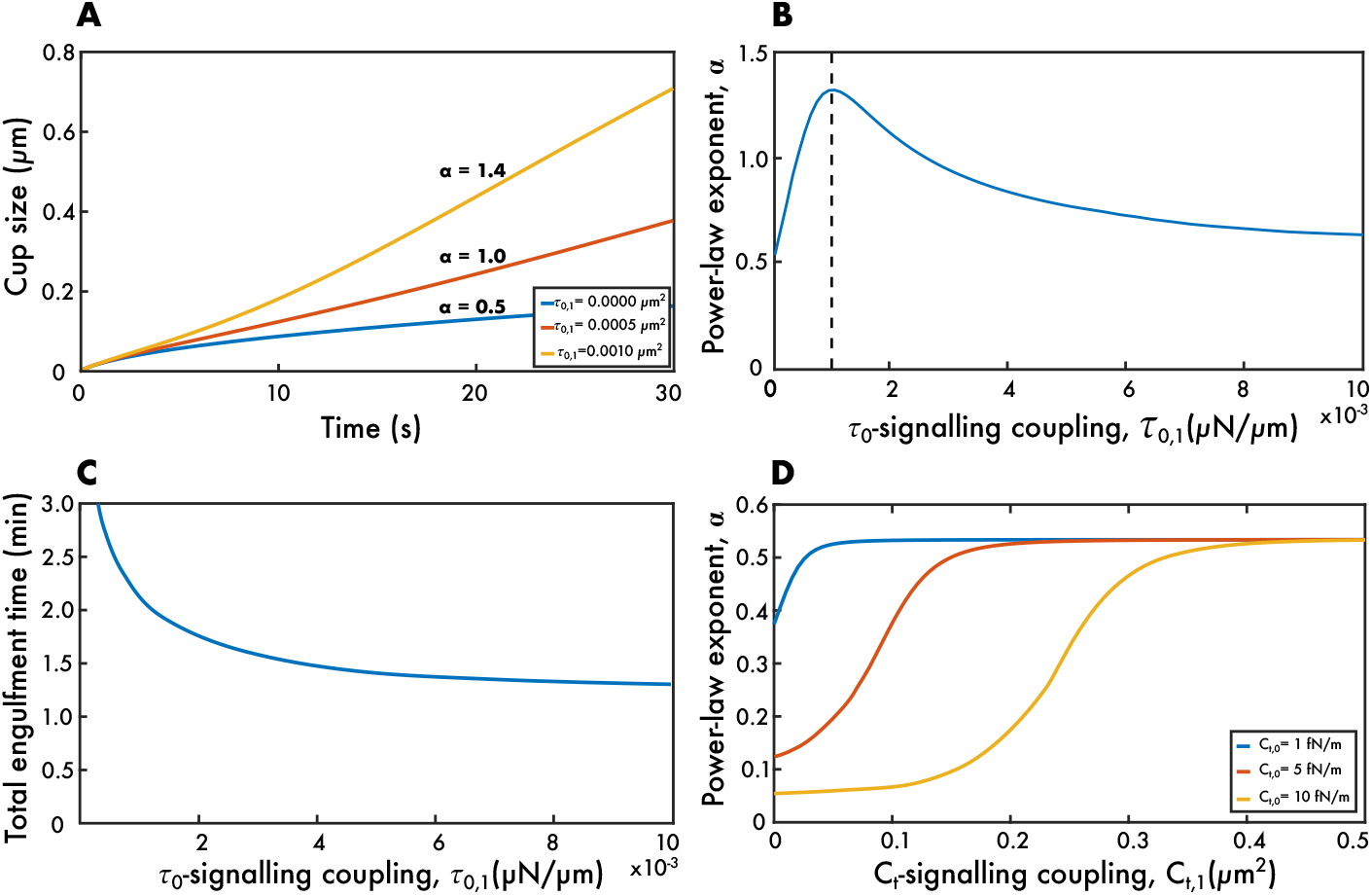
The effect of including signalling in the model. (A-C) The results of coupling *τ*_0_ to the signalling molecule. (A) The cup size *a* against time *t* shows that signalling can increase the cup growth behaviour to be quicker than 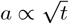. By changing the coupling constant *τ*_0,1_, it is possible to obtain linear or even super-linear growth in time. (B) The growth behaviour *α* (found by fitting *a* = *Bt*^*α*^) starts around 0.5 for no signalling, increases to a maximum as *τ*_0,1_ is increased (dashed line) and then decreases back towards 0.5 as *τ*_0,1_ is further increased. (C) The total engulfment time decreases as *τ*_0,1_ increases, asymptoting to a constant value corresponding to the case where *ρ*_+_ = 0. (D) The results of instead coupling *C*_*t*_ to the signalling molecule, showing that *α* increases as *C*_*t*,1_ increases, but never exceeds 0.5. Parameter values for panels (A-C): *D* = 1*µ*m^2^*/*s, *C*_*c*_ = 10, *C*_*b*_ = 15, *C*_*t*_ = 10^−4^fN*/*m, *τ*_0,0_ = 0.014fN*/*m, *k*_*B*_*T* = 4 × 10^−21^J, *ρ*_*L*_ = 500*µ*m^−2^, *ρ*_0_ = 50*µ*m^−2^, *R*_target_ = 2*µ*m, *R*_cell_ = 10*µ*m, *D*_*S*_ = 1*µ*m^2^*/*s, *β* = 0.5s^−1^, *η* = 0.5s. Parameter values for panel (D) as for panels (A-C) but with *C*_*t*_ varying and *τ*_0,1_ = 0.

Biologically, it is quite plausible that signalling could affect the membrane properties, particularly its tension, in order to drive efficient engulfment. For instance, small GTPases and PI3K are known to modulate actin polymerisation and membrane tension, both of which are essential for cup extension [49, 50]. Our model predicts that increasing signalling can accelerate phagocytosis by decreasing local membrane tension. This aligns with experimental observations where inhibition of signalling pathways leads to stalled cups [51]. Furthermore, the emergence of switch-like engulfment dynamics at high Hill coefficients may reflect a biological mechanism for target discrimination, allowing phagocytes to commit rapidly to opsonised or damaged cells while ignoring weaker, non-pathogenic stimuli [17, 52].

### Full model behaviour

Finally, we examine how the full model (including both membrane tension and signalling coupled to *τ*_0_) depends on the various parameters. To this end, we individually vary each of the model parameters and plot how each affects the total engulfment time *T* (see Fig. 5).

**Fig 5.**
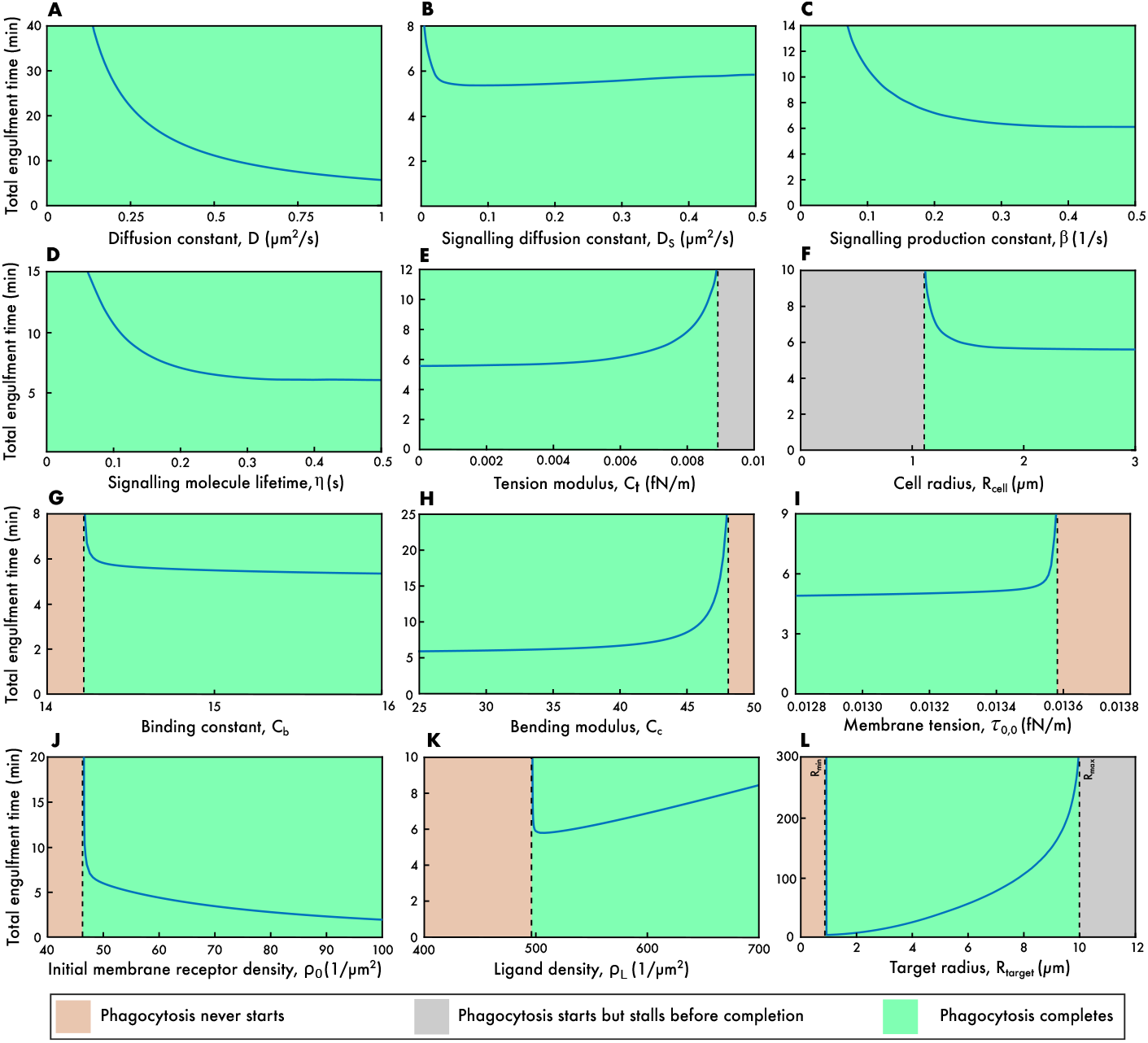
Effect of model parameters on the total engulfment time. (A) The receptor diffusion constant, *D*. (B) The signalling molecule diffusion constant, *D*_*S*_. (C) The signalling molecule production constant, *β*. (D) The signalling molecule lifetime, *η*. (E) The tension coefficient, *C*_*t*_. (F) The cell radius, *R*_cell_. (G) The binding affinity, *C*_*b*_. (H) The curvature constant, *C*_*c*_. (I) The initial membrane tension per unit length, *τ*_0,0_. (J) The initial receptor density, *ρ*_0_. (K) The target ligand density, *ρ*_*L*_. (L) The target radius, *R*_target_. Default parameters used in these simulations: *D* = 1*µ*m^2^*/*s, *C*_*c*_ = 10, *C*_*b*_ = 15, *C*_*t*_ = 10^−4^fN*/*m, *τ*_0,0_ = 0.014fN*/*m, *τ*_0,1_ = 0.005*µ*N*/µ*m, *k*_*B*_*T* = 4 × 10^−21^J, *ρ*_*L*_ = 500*µ*m^−2^, *ρ*_0_ = 50*µ*m^−2^, *R*_target_ = 2*µ*m, *R*_cell_ = 10*µ*m, *D*_*S*_ = 1*µ*m^2^*/*s, *β* = 0.5s^−1^, *η* = 0.5s.

Each parameter choice leads to one of three possible outcomes—phagocytosis fails to start, phagocytosis starts but stalls before completion, and phagocytosis completes—with only the last case having an associated total engulfment time. We find that our various model parameters separate into four distinct categories based on how the outcome of phagocytosis depends on the parameter.

The first category, which includes the receptor diffusion constant and the three signalling molecule constants, are the cases where engulfment always completes for all values of the parameter (see Fig. 5(A)-(D)). In the case of the receptor diffusion constant *D*, increasing values of *D* lead to quicker engulfment with *T* tending to zero for very large *D* (Fig. 5(A)). This behaviour is very close to *T* ∝ 1*/D*, which is the exact analytic result in the case that signalling is turned off (*i*.*e*. when *τ*_0,1_ = 0). This matches the work of Jaumouillé and Waterman, who described how dynamic receptor redistribution, including lateral diffusion in the membrane, is crucial for assembling a functional phagocytic cup [53]. Supporting this further, Zhang *et al*. showed that receptor mobility enables Fc receptors to cluster efficiently at the site of contact, which is necessary for the cell to commit to phagocytosis [54].

The signalling production constant *β* (Fig. 5(C)) and signalling lifetime *η* (Fig. 5(D)) have similar behaviour in that increased *β* or *η* lead to quicker engulfment. However, unlike *D*, both *β* and *η* asymptote to a non-zero value of *T* for large values. This is because very large levels of signalling (and hence large values of *S*_+_) effectively set *τ*_0_ to zero, but this still leaves a non-zero right-hand side to Eq. (7).

The final parameter in this first category is the signalling molecule diffusion constant, *D*_*S*_, which shows interesting non-monotonic behaviour, with an intermediate *D*_*S*_ corresponding to the quickest engulfment (Fig. 5(B)). Low values of *D*_*S*_ result in the signalling molecule concentrated near the base of the cup (where *r* = 0), leading to small values of *S*_+_ and so slow engulfment. Conversely large *D*_*S*_ results in very quick spreading out of *S* over the whole membrane, which again results in relatively small *S*_+_ and so small *T*. This of course assumes that the effect of the signalling molecule is only felt at the edge of the phagocytic cup (*i*.*e*. at *r* = *a*), an assumption that is unlikely to be biologically accurate and that could be relaxed in future models.

The second category of behaviour are those parameters that show a switch between complete and stalled engulfment. Phagocytosis starts for all parameter values, but sometimes stalls before complete engulfment. This can only occur due to the presence of the *a*(*t*)^2^ term in Eq. (7), and so is seen only for the tension coefficient *C*_*t*_ and the cell radius *R*_cell_. For the case of *C*_*t*_, phagocytosis completes for small values, but stalls for *C*_*t*_ above some critical value when the membrane tension builds up during engulfment to such an extent that further cup growth become impossible (Fig. 5(E)). For the case of *R*_cell_ the opposite behaviour is seen (Fig. 5(F)): large cells can engulf the target, but lack of available spare membrane in smaller cells results in incomplete phagocytosis. Experimental studies agree with this and show that the phagocytic capacity of a cell is constrained by its size [44].

The third category are the parameters that switch from phagocytosis never starting to phagocytosis completing. There is no intermediate stalled regime, just a sharp switch between no engulfment at all and complete target wrapping (Fig. 5(G)-(K)). These cases are related to the parameters in Eq. (7) that are not part of the *a*(*t*)^2^ term. If *ρ*_+_ ≥ *ρ*_0_ at the start of engulfment (*i*.*e*. when *a* = 0) then engulfment never starts, otherwise phagocytosis always completes.

There are five parameters in this category: the binding constant *C*_*b*_, bending modulus *C*_*c*_, initial membrane tension *τ*_0,0_, initial membrane receptor density *ρ*_0_ and ligand density *ρ*_*L*_. For the initial membrane receptor density, failed phagocytosis is associated with low values of *τ*_0,0_ due to there being no inward flux of receptors at the edge of the cup (Fig. 5(J)). Similarly, small binding constants do not show even partial engulfment since the energy gain from receptor-ligand binding is insufficient to drive engulfment (Fig. 5(G)). This agrees well with the work of Freeman and Grinstein who emphasised that strong receptor-ligand interactions help stabilise the contact zone and amplify signalling [7].

The bending modulus and initial membrane tension display the opposite behaviour: it is large values of *C*_*c*_ (corresponding to high energy penalty from bending the membrane around the target; Fig. 5(H)) and large values of *τ*_0,0_ (Fig. 5(I)) that are associated with failure to phagocytose. This idea is supported by Herant *et al*., who used mechanical modelling and live-cell imaging to show that neutrophils must lower their membrane tension to allow for the large deformations required during engulfment [15]. Similarly, Hallett and Dewitt gave evidence that flexible membranes are essential for efficient phagocytic cup formation, especially in neutrophils [55].

The final parameter in this category is the ligand density *ρ*_*L*_, where the total engulfment time displays interesting non-monotonic dependence, with the quickest engulfment occurring for an intermediate *ρ*_*L*_ (Fig. 5(K)). Small values of *ρ*_*L*_ do not lead to any engulfment since there is then relatively little energy gain from receptor-ligand binding. Conversely, large values of *ρ*_*L*_ do lead to complete engulfment but this takes a long time due to the high density of ligands on the target that must be bound. This is consistent with earlier energy-based models of endocytosis, which showed that there is an optimum ligand density where membrane wrapping is energetically most favourable. Specifically, Gao *et al*. [33] and Shadmani *et al*. [23] demonstrated that particle uptake is optimised at intermediate ligand densities due to a balance between adhesive forces and membrane deformation energy. Richards and Endres reported the same results for phagocytosis and concluded that excessive ligand coverage can lead to slower uptake [17].

It is worth noting that the lack of a stall region in the third category of parameters is related to the size of the initial membrane tension, *τ*_0_. Once engulfment starts there are two competing factors determining whether phagocytosis completes: signalling (which increases the rate of engulfment) and the membrane tension (which slows engulfment). With our parameter values, signalling is always the dominant effect.

However, for other parameter choices (*e*.*g*. small *τ*_0,1_) this is no longer the case and stalling can occur, albeit with unrealistically long engulfment times when phagocytosis does complete. In such cases, the third category would display three regions (similar to Fig. 3D for *R*_target_ *<* 2*µ*m), with a no-wrapping region transitioning into a stall region and then into a complete-engulfment region as the relevant parameter is varied.

The final category of behaviour applies only to the target radius, *R*_target_ (Fig. 5(L)). This is similar to Fig. 3D, but now for the full model. However, there are some important differences. First, there is no longer an initial stall region just below *R*_min_. This is for exactly the same reason as for the third category above: signalling dominates over membrane tension for our parameter choices so that *ρ*_+_ always decreases during engulfment. Also as above, this stall region can be reintroduced by choosing unrealistic parameters for *τ*_0_, with the plot then looking similar to Fig. 3D.

Second, the quickest engulfment time is now around 120 s (cf. 90 mins in the case without signalling), demonstrating that our full model correctly captures the typical engulfment times seen experimentally. Notably, the critical target radius that corresponds to the shortest engulfment time is similar between the full model and no-signalling model, around *R*_target_ = 0.9*µ*m in both cases, consistent with previous models [17, 23, 33].

Third, the maximum size target that can be engulfed increases from around 5*µ*m in the no-signalling model to around 10*µ*m here, which again agrees with experimentally-observed values for *R*_max_ [15]. This result highlights the critical role of actin-mediated force generation in enabling efficient engulfment, particularly for larger targets. As noted by van Zon *et al*. [18], actin polymerisation not only provides the force necessary to deform the membrane around a target, but also actively drives cup progression and membrane advancement. Without actin involvement, phagocytosis either does not complete or is prohibitively slow, especially for target sizes beyond those of a typical bacterium.

## Conclusions

In this paper, we have developed a model of phagocytosis that improves upon our previous model by incorporating the effects of membrane tension. Our findings highlight the critical role played by tension during phagocytosis and demonstrate how it can significantly affect the engulfment dynamics. One of our key results is that, when membrane tension is considered, phagocytosis can began but then stall before completion, a phenomenon that was not possible in our previous model but that has been observed experimentally. This agrees well with the theoretical results of van Zon *et al*. [18] who found, using a different model, that engulfment often stalls around half way.

We also included a signalling component to our model, where bound receptors cause altered engulfment dynamics. This is important during phagocytosis (the engulfment of relatively large *>* 0.5*µ*m particles), which requires active processes, typically related to actin, in order to be successful. By coupling signalling to our new addition of membrane tension (either via the parameters *τ*_0_ or *C*_*t*_) we found that the total engulfment time can be substantially reduced down to a couple of minutes, agreeing well with experimentally-measured values.

In addition, the inclusion of signalling allows much larger targets to be engulfed (10*µ*m in radius rather than 5*µ*m in the no-signalling model), which again matches well with experimental results, where the maximum size that can be phagocytosed has been shown to be about 20*µ*m in diameter [15].

We also showed that signalling can fundamentally alter the cup growth behaviour, away from the 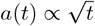 predicted from non-signalling models. In fact, just by coupling signalling to membrane tension it is possible to achieve linear cup growth or even sup-linear growth. Previous models could achieve this only be including a drift component to the receptor dynamics.

Finally, we examined how our full model depends on the model parameters. As in our previous model, we found interesting dependence on the target size *R*_target_. We predict that targets that are too small (*R*_target_ *<* 1*µ*m) or too large (*R*_target_ *>* 10*µ*m) cannot be engulfed via phagocytosis, although other types of endocytosis are able to ingest smaller targets. In addition, although we find that the total engulfment time *T* is non-monotonic with *R*_target_, the region with *T* decreasing is now so small that it can effectively be ignored: such a region is unlikely to be experimentally observable.

As with any model, we have made a number of assumptions and simplifications that are worth bearing in mind. First, we have assumed that growth of the phagocytic cup is chiefly controlled by the motion of receptors coupled to a single, proxy signalling molecule. The fact that we obtain realistic engulfment dynamics justifies this approach. Second, we assumed that engulfment is circularly symmetric when viewed from directly above. This is not the case in general, but leads to a substantially simpler, one-dimensional system that is tractable analytically and numerically. Third, we ignored any system noise, which is undoubtedly present in any real cell. This again leads to a simpler, easier-to-understand system. Fourth, we assumed that all receptor-ligand bonds are stable and irreversible, resulting in zipper-like engulfment [16]. However, recent biophysical studies have suggested that receptor-ligand interactions are in fact transient [56], with their kinetics influenced by the mechanical forces generated during membrane deformation [57].

Future work may aim to relax some of these model assumptions by, for example, considering a more complete three-dimensional system that includes noise. In addition, future versions of the model could include a dynamic binding framework, where receptors are able to unbind from ligands with a rate depending on the mechanical force along the cup and membrane curvature.

Another possible future direction is to consider that only a portion of the available ligands on the target need to be bound by receptors. Experiments have shown that both the density and spatial arrangement of ligands can significantly impact phagocytic efficiency [54, 58]. In addition, in real biological systems, not all ligands may be accessible, with factors like blockage by other proteins preventing full receptor binding. Incorporating a variable ligand-binding fraction into the model, possibly influenced by properties of the membrane or patterns of receptor clustering, could be a useful addition [59].

Finally, we here assumed that the role of signalling is to modulate one of the membrane tension parameters, either *τ*_0_ or *C*_*t*_. Future work could consider the possibility that both parameters are simultaneously affected. Further, the signalling density *S* could couple to entirely different factors, such as the membrane bending modulus *C*_*c*_ (with signalling lowering the modulus) or the receptor-ligand bond strength *C*_*b*_ (with signalling increases the bond strength). However, although these options would capture different biology, it is unlikely that they would substantially alter our conclusions on how signalling affects engulfment dynamics.

By including membrane tension in our previous model, we have developed a more realistic model of phagocytosis that demonstrates a richer set of behaviours. Continuing along this path in future, with the addition of further biophysical components (such as the role of the cytoskeleton and other intracellular components), will help elucidate the underlying mechanisms that govern phagocytosis, including its vital role in the immune system, with the ultimate aim of a complete understanding of one of nature’s most complex processes.

## Methods

### Numerical solution

We numerically solve our model system between *r* = 0 and *r* = *L* using the Euler method with Δ*t* = 2.5 × 10^−4^s, Δ*r* = 0.01*µ*m and *L* = 50*µ*m. Finite difference versions of Eqs. (1), (2) and (8) are used at each time step. For maximum accuracy, *a*(*t*) is allowed to vary continuously (*i*.*e*. it is not restricted to lie on the lattice). The initial and boundary conditions are summarised in Table 1, although we initialise the system with a small perturbation away from these values in order to trigger engulfment. We verified that reducing Δ*r* or Δ*t* has negligible effect. Further details are given in our previous work [17, 22].

### Parameter values

Various bending constants have been measured in the literature, with values depending on cell type and ranging from 1.5 *×* 10^−21^J to 1.5 *×* 10^−19^J [39, 60]. Given that *k*_*B*_*T* = 4 *×* 10^−21^J, we choose an intermediate value of *C*_*c*_ = 10. The binding constant is taken as *C*_*b*_ = 15, which is the Fc*γ*R-IgG binding free energy (15*k*_*B*_*T*) [61]. Based on the tension model we introduced here and by fitting to the measured engulfment times and range of target sizes that can be engulfed, we use *C*_*t*_ = 10^−4^fN*/*m, *τ*_0,0_ = 0.014fN*/*m and *τ*_0,1_ = 0.005*µ*N*/µ*m. Typical literature values are used for the initial receptor density *ρ*_0_ = 50*µ*m^−2^ [33] and the ligand density *ρ*_*L*_ = 500*µ*m^−2^ [62]. The receptor diffusion constant and signalling parameters are chosen as in our previous studies with *D* = *D*_*S*_ = 1*µ*m^2^*/*s, *β* = 0.5s^−1^ and *η* = 0.5s [17, 22].

## Funding information

DMR gratefully acknowledges financial support from the Medical Research Council (MR/P022405/1), and from the Biotechnology and Biological Sciences Research Council and the National Centre for the Replacement, Refinement and Reduction of Animals in Research (NC/X002268/1). PS thanks the National Centre for the Replacement, Refinement and Reduction of Animals in Research for funding via an Early Career Engagement award (NC/ECE0028/1). The funders had no role in study design, data collection and analysis, decision to publish, or preparation of the manuscript.

## Notes

### Competing Interest Statement

The authors have declared no competing interest.

